# Cell position is the primary determinant of cell identity in the uniseriate filament of the brown alga *Ectocarpus*

**DOI:** 10.64898/2026.01.21.700896

**Authors:** Denis Saint-Marcoux, Bernard Billoud, Sabine Chenivesse, Carole Duchêne, Aude Le Bail, Jane A. Langdale, Bénédicte Charrier

**Author notes:** These authors contributed equally to this work (Denis Saint-Marcoux, Bernard Billoud).

## Abstract

The brown alga *Ectocarpus* is a morphologically simple organism, with cells in the growing filament changing shape and position over time. In this study, we investigated the roles of cell shape, position, and age in establishing cell identity in *Ectocarpus*. Using laser capture microdissection, we isolated cells with different relative ages, shapes and positions from young sporophytes, and analysed their transcriptomes using differential RNA-seq to attribute a molecular signature to each cell. Five distinct cell transcriptomic signatures were found along the filament of WT *Ectocarpus*; the apical cell signature differed most from the others. To dissociate cell shape, age, and position, we analysed the transcriptomes of two mutants in which the relationships between these three parameters were disrupted. We found that the transcriptomic profile observed in the WT was significantly altered in these mutants. Overall, our data revealed that molecular cell signatures depend primarily on cell position along the filament and secondarily on cell shape.

## Introduction

Cell differentiation is the process by which cells acquire the characteristics that distinguish them from other cells, thus establishing their cell type or identity. The characteristics that define cell identity include shape, age (in the case of progenitor cells, for example), behaviour (such as growth rate and metabolic activity), and gene expression (Chen et al., 2015; Shainer et al., 2025). In both animals and plants, cell identity may be influenced by extrinsic signals, such as the environment, for example the positional information (Cole et al., 2024; Rusnak et al., 2024) that can be imparted from neighbouring cells through biophysical (Mihajlović and Bruce, 2017; Vermeer et al., 2014) or biochemical signals (Winnicki, 2020). It can also respond to intrinsic signals, such as cell lineage, but also cell shape (Chen et al., 2020; Haftbaradaran Esfahani et al., 2019; Luxenburg and Zaidel-Bar, 2019; Sablowski and Gutierrez, 2022) and age (da Silva and Schumacher, 2021; Lans and Hoeijmakers, 2006).

These factors are closely interconnected making it difficult to unpick their individual contributions. The shape and age of a cell type can sometimes be manipulated physically or genetically (see (Kordyum et al., 2019; Zegman et al., 2015; Zhang et al., 2005), but altering cell position without massively disturbing the system is not possible, particularly in organisms with walled cells like plants, fungi and algae.

In organisms composed of uniseriate filaments, which are one cell wide, all cells except the apical cells are surrounded by only two neighbours. Relative cell age can also easily be inferred when growth is restricted to the apical cells. Therefore, uniseriate filamentous organisms are valuable models for distinguishing the effects of cell shape, relative age and position on cell differentiation. Although the hyphae of fungi, the protonemata of mosses, and the filaments of algae satisfy most of these criteria, cell differentiation usually cannot be tracked based on a change in cell shape because the cell morphology remains mainly unchanged along the filament. In the filamentous brown alga *Ectocarpus*, by contrast, cell shape changes with both position and relative age. Since publication of its genome sequence (Cock et al., 2010), this alga has pioneered molecular studies of brown algae development and morphogenesis (Arun et al., 2019; Billoud et al., 2014; Billoud et al., 2015; Coelho, 2024; Coelho et al., 2020; Godfroy et al., 2017). Notably, this is an excellent model for tracking cell differentiation based on changes in cell shape as apical growth proceeds (Billoud et al., 2008; Charrier et al., 2008; Le Bail et al., 2008; Le Bail et al., 2011). This morphology prompted us to attempt to disentangle the contributions of cell shape, age and position to cell differentiation in *Ectocarpus* by comparing the transcriptomes of specific cell types in wild type and mutant strains. To analyse the transcriptomes of specific cell types, we used laser capture microdissection, which unlike cell dissociation single cell (sc)RNA-seq methods, retains cell positional information without causing stress and impacting the transcriptional state of the cell (Espina et al., 2006; Kubo et al., 2019). Our results show that cell position is the primary determinant of cell identity in this organism.

## Results

### Wild-type *Ectocarpus* sporophytes comprise five cell types with distinct shapes, positions and relative ages

When mature, *Ectocarpus* is a multicellular brown alga (Ectocarpales, Phaeophyceae) composed of branched filaments (Fig. 1A). This structure develops from a filament comprising a single chain of cells (Fig. 1B). The morphogenesis of this filament and the definition of the different cell morphotypes have been described previously (Billoud et al., 2008; Charrier et al., 2021; Le Bail et al., 2008; Le Bail et al., 2011; Nehr et al., 2011). To study the role of cell position, shape and age in determining cell identity in *Ectocarpus*, we allocated a specific coordinate to each cell based on its position (Greek letter), shape (capital Roman letter) and relative age (number; Fig. 1C). Upon germination, the zygote produces a first apical cell (position ɑ) that elongates and divides transversely, thus producing all the cells of the filament. Based on this growth mode, relative cell age was inferred from the cell division rate of the apical cell and the ranked position of its daughter cells (see Material and Methods for a detailed explanation of the definition of relative age). The apical cell, present at each end of the filament, is always the youngest cell (defined as estimated age 1). It has an elongated cylindrical shape (shape E), which we measured as having a length/width ratio (L/w) ≥ 1.5. Successive transverse divisions of the apical cell, αE1, result in formation of sub-apical (position β), elongated (shape E) cells that are older (age 2). As the filament grows, βE2 cells progressively become rounder and may divide once. Intermediate (shape I) cells with a L/w ratio included in [1.1: 1.5[ are older (age 3) and located in a more central position (γ) than the βE2 cells. In turn, these γI3 cells eventually differentiate into fully round cells (L/w ratio ɛ [1.0: 1.1[) that occupy a central position (position δ) in the filament and are older (age 4). In this species, branching can occur from all cells except the apical cell αE1 (Le Bail et al., 2008; Nehr et al., 2011). However, in this study, to standardise positional context at the 10-day stage (∼15–20-cell filaments), the branching samples were restricted to branched δR4 cells, which were named δB4 cells (B for branching). By using bright field microscopy to monitor cell differentiation over 7 days, we observed that the transition from αE1 to δR4 cell occurs in ∼ 4 days in mature *Ectocarpus* sporophytes (Movie 1; Fig. S1A). During this time, the length of the δR4 cells decreased by 36.9% when compared with βE2 cells, their width increased by 44% and their volume increased fourfold (Fig. S1B).

**Figure 1.**
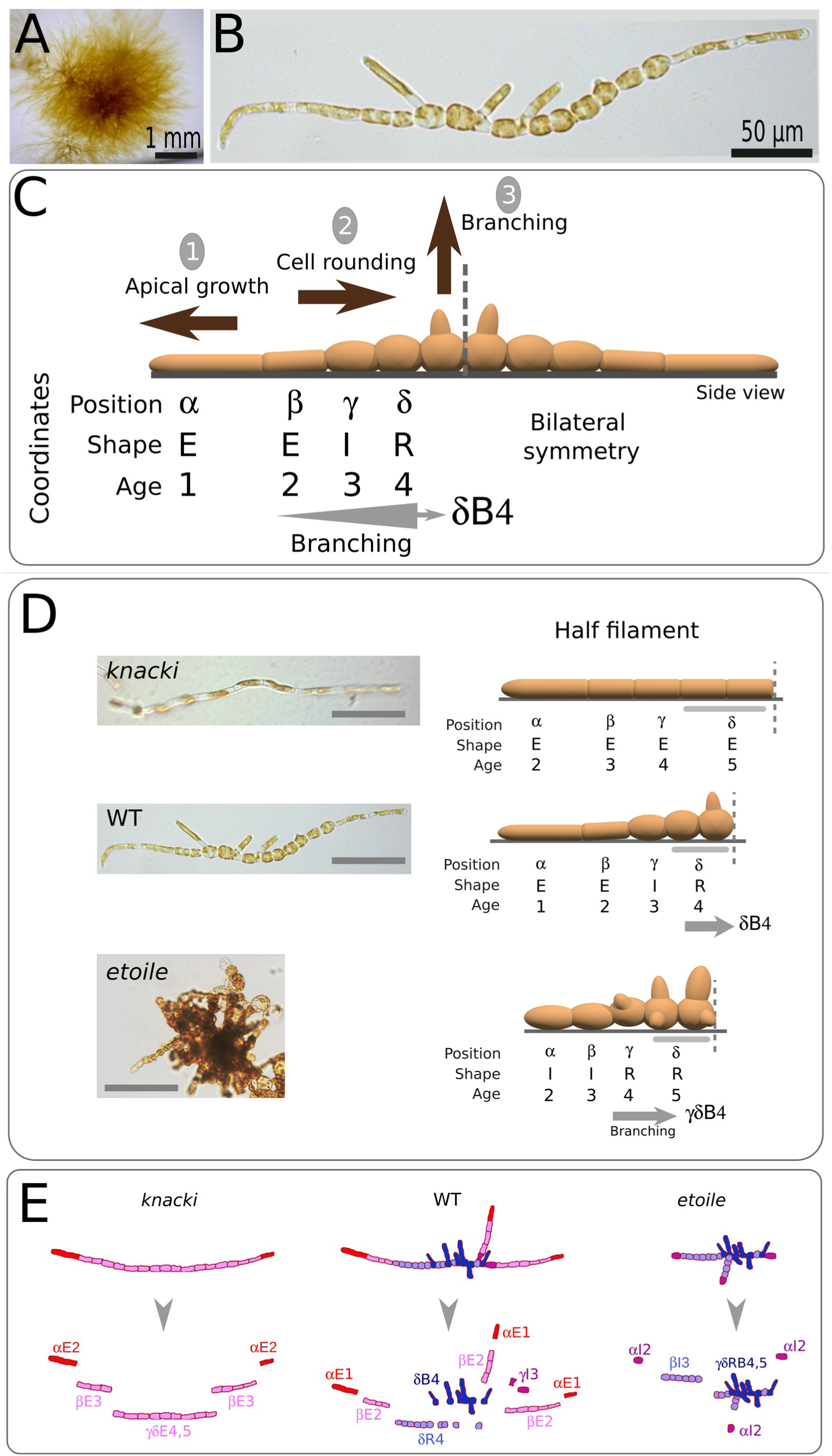
: Morphology of early *Ectocarpus* WT and mutant sporophytes and cell types. In the WT, *Ectocarpus s*p. sporophyte filaments display five cell types based on their shape, location and age. **(A)** One month-old *Ectocarpus* sporophyte made of branched filaments. **(B)** 10-day old *Ectocarpus* sporophyte filament showing cylindrical cells at the distal end and spherical cells in the more central region, together with primary branches. **(C)** Schematised representation of an early sporophytic filament. Three main cellular processes take place during growth: 1) apical growth, 2) cell rounding and 3) branching. Three cell types were defined along the filament, from the tip to the center, according to their shape (E, I and R, based on the width/length ratio), position (α, β, γ and δ) and age (1, 2, 3 and 4; defined as the time since their formation from the apical cell division; see the Material and Methods section for a detailed explanation). Branching occurs on all cell types except the apical cell. In the context of this study, only branching taking place on δR4 was considered. Cell differentiation takes place symmetrically from both ends. **(D)** Overall morphology of the mutants *etl* and *kna*, as compared to WT. Photos of ∼ 2-week old organism are shown, together with schematised half filaments. Note that in the WT, cell types are represented by only one cell of each type, but there are usually several cells of each type grouped together (except the apical cell, which is always single by definition). Scale bar = 50µm. Cell types for the WT and for each mutant are characterised by (i) position relative to the global filament organisation, (ii) cell shape computed as the length/width ratio and (iii) cell age since division from the apical cell. **(E)** Laser ablation method, for sporophyte filaments of WT (center), and *kna* (left) and *etl* (right) mutants. Each cell type in the WT is represented with a different color : Red for the apical cell type (α), pink for sub-apical (β), purple for transitory intermediate cell stage (γ) in which cell shifts from cylindrical to spherical cell shape, light blue for central cells (δ) and dark blue for B (branched cells located mostly in δ regions). Mutants *etl* and *kna* are depleted in specific cell shapes: E in *etl*, I and R in *kna.* In these mutants, cell age 1 is missing, cell age 5 is gained.

In summary, because of apical cell elongation and division, the cells at the tip of the *Ectocarpus* filament are the youngest and most polarised. The most central cells are the oldest and most isotropic, as they result from the progressive rounding of the apical cell’s daughter cells. Thus, in the wild-type (WT) *Ectocarpus* sporophyte, cell age, shape, and position are intrinsically linked, making it difficult to attribute the relative contributions of each of these three factors to the acquisition of cellular identity.

### The relationship between cell shape, age and position is perturbed in *knacki* and ***etoile* mutants**

To analyse the relative contributions of cell shape, position and age to cell identity, we used UV mutagenesis to produce two cell shape mutants, *knacki (kna)* and *etoile (etl)*. In *kna* filaments, cells in the γ position did not differentiate into I-shaped cells, but remained E-shaped (Fig. 1D; Fig. S2A). Consequently, R cells were absent from the δ position. Thus, the *kna* filament is composed only of E-shaped cells that are similar in size to WT cells (length 23.5 µm for 25.8 µm in the WT; width 10 *vs* 10.5 µm; T-test P-value > 0.05; Fig. S2A). Time-lapse measurements showed that *kna* mutant filaments grew more slowly than the WT (1.81 µm.h^−1^ *vs* 2.54 µm.h^−1^; Fig. S2C). We reasoned that cells in a filament with a slower growth rate are older than those in faster-growing filaments in the same relative position (see Materials and Methods for a detailed explanation). Thus, *kna* filaments, which grow more slowly than WT filaments and are composed of only E-shaped cells, are formed successively from the apex of αE2, βE3, γE4 and δE5 cells (Fig. S2D). In this mutant, therefore, cell shape is not associated with cell age or position. The relationship between cell age and position differs between the *kna* mutant and the WT.

The phenotype of the *etl* mutant was the converse of that of *kna* (Fig. 1D). The *etl* mutant filaments did not form E-shaped cells and both the α cell – which is involved in tip growth as in the WT – and the β cell, were I-shaped. The other cells of the filament were R-shaped, and slightly larger than WT R-shaped cells (cell diameter of 24.6 *vs* 20.5 µm in the WT, as reported in (Le Bail et al., 2011); Fig. S2B). Thus, *etl* is made only of I- and R-shaped cells. Furthermore, apical growth was slower in *etl* filaments and eventually stopped after about 7 days (Le Bail et al., 2011). Time-lapse measurements showed that *etl* mutant filaments grew more slowly than the WT (1.54 µm.h^−1^ *vs* 2.54 µm.h^−1^; Fig. S2C). Consequently, in *etl* filaments, the apical and its descendants are older than their WT counterparts (Fig. S2D). Therefore, we characterised the *etl* filament as successively composed of αI2, βI3, γR4 and δR5 (see Fig. 1D). Thus, in this mutant both cell shape and age differ in their relationship to cell position when compared with the WT.

Together, therefore, the two mutants and the WT present a range of morphological and growth parameters that include variations in the rate of cell shape changes, the dynamics of cell growth, impacting cell age, and the overall spatial organisation. Thus, they provide a genetic system to explore which of these factors contributes most to cell identity at the molecular level.

### WT cell types show major transcriptomic differences

To determine the molecular identities of the five cell types identified in WT filaments, we manually isolated cells of the same type using laser capture microdissection, which allows spatial information to be conserved (Shapiro et al., 2013). Cells αE1, βE2, γI3, δR4 and δB4 were dissected from at least 50 two-week-old WT filaments containing ∼50 cells (Fig. 1E). Apart from the αE1 cell type, which is present at each end of the filament, each of the other cell types contains two to four adjacent cells within the filament. To avoid losing cell material due to the thickness of the laser beam (Shapiro et al., 2013), we isolated adjacent cell-types from separate filaments. We extracted and sequenced the RNA from the isolated cells. Thirty million reads were obtained for each cell type, summed from three independent biological replicates (∼ 10^7^ reads each).

Before analysing the transcription profiles in different cell types, we assessed the extent to which our captured transcriptome represents overall gene expression in *Ectocarpus*. The sporophytic phase lasts as long as the gametophytic phase (i.e. about six weeks; (Charrier et al., 2008). Although 10-day-old sporophytic filaments represent just 12% of the total life cycle duration, they express up to 77.7% (12,999 transcripts) of the predicted total gene number (16,738). Furthermore, we found that 80% of genes were sporophyte-biased and 50% were sporophyte-specific (Table S1; (Lipinska et al., 2019). More surprisingly, 45% of gametophyte-biased genes and at least 10% of gametophyte-specific genes were also expressed in early sporophytes. This suggests that most genes are expressed throughout both life cycle generations, as has previously been observed in brown algae (Ratchinski et al., 2025). Overall, our dataset represents over 97% of the predicted transcriptome.

Each of the five cell types expressed about 77% of the total number of predicted genes. This suggests that a significant proportion of the transcriptome was expressed in all cell types. We also found 1596 genes that were significantly differentially expressed in at least one pairwise comparison of transcript abundance in αE1, βE2, γI3 and δR4 cells (p-value < 0.05; Table S2). Pairwise comparisons between adjacent cell types showed a progressive decrease in the number of differentially expressed genes from the tip to the centre of the filament (Table 1): 472 genes were differentially expressed between αE1 and βE2; 218 between βE2 and γI3, and only 7 between γI3 and δR4. Therefore, the transitions from αE1 to βE2 and from βE2 to γI3 encompassed most of the transcriptional changes observed along the filament during cell differentiation. This suggests a significant reconfiguration of gene expression accompanies the change from αE1 to βE2, and a more minor reconfiguration occurs during the rounding process from βE2 to δR4 as gene expression becomes progressively more stable. We saw little difference in gene expression between γI3 and δR4, making these two cell types virtually identical in terms of their transcriptomes.

**Table 1:** Number of genes showing differential expression (FDR < 0.05) in all pairwise comparisons.

| Cell type | αE1 | βE2 | γI3 | δR4 |
| --- | --- | --- | --- | --- |
| αE1 | | | | |
| βE2 | 472 | | | |
| γI3 | 1024 | 218 | | |
| δR4 | 1321 | 496 | 7 | |

These transcriptomics data reflect the dynamics of WT cell differentiation in time and space, at a spatial resolution of one cell (∼ 15 µm) and a temporal window of ∼ 1 day (corresponding to growth of ∼ 60 µm, i.e. 1–2 cells at each end of the filament; (Rabillé et al., 2019a). Also, they show that gradual changes in cell identity defined by shape, position and age, are accompanied by progressive modifications of their transcriptome signatures.

### Apical cell identity is distinct from the other cell types

To identify the major biological processes associated with differentiation from the αE1 to the δR4 cell, we clustered differentially expressed genes based on the direction of change in the three cell types αE1, βE2 and γI3 when compared to δR4. The δR4 cell was used as a reference because it is the final state of cell differentiation in the filament. We found that most genes were grouped into four main clusters: ‘b’ (244 genes), ‘c’ (493 genes), ‘j’ (407 genes) and ‘k’ (171 genes); Fig. 2A). The remaining genes were mainly grouped into two additional clusters: ‘d’ (37 genes) and ‘f’ (also 37 genes). Clusters ‘b’ and ‘c’ comprised genes whose activity increased during cell differentiation from αE1 to δR4, whereas clusters ‘j’ and ‘k’ comprised genes whose expression decreased during differentiation from αE1 to δR4. In clusters ‘j’ and ‘k’, a notable drop in transcript levels between αE1 and βE2 was even observed. In clusters ‘d’ and ‘f’, a tipping point was observed where the level of transcripts increased or decreased, centred on βE2. This resulted in the isolation of αE1 from the other cell types and suggests that the apical cell has a molecular identity distinct from all other cells of the filament.

**Figure 2:**
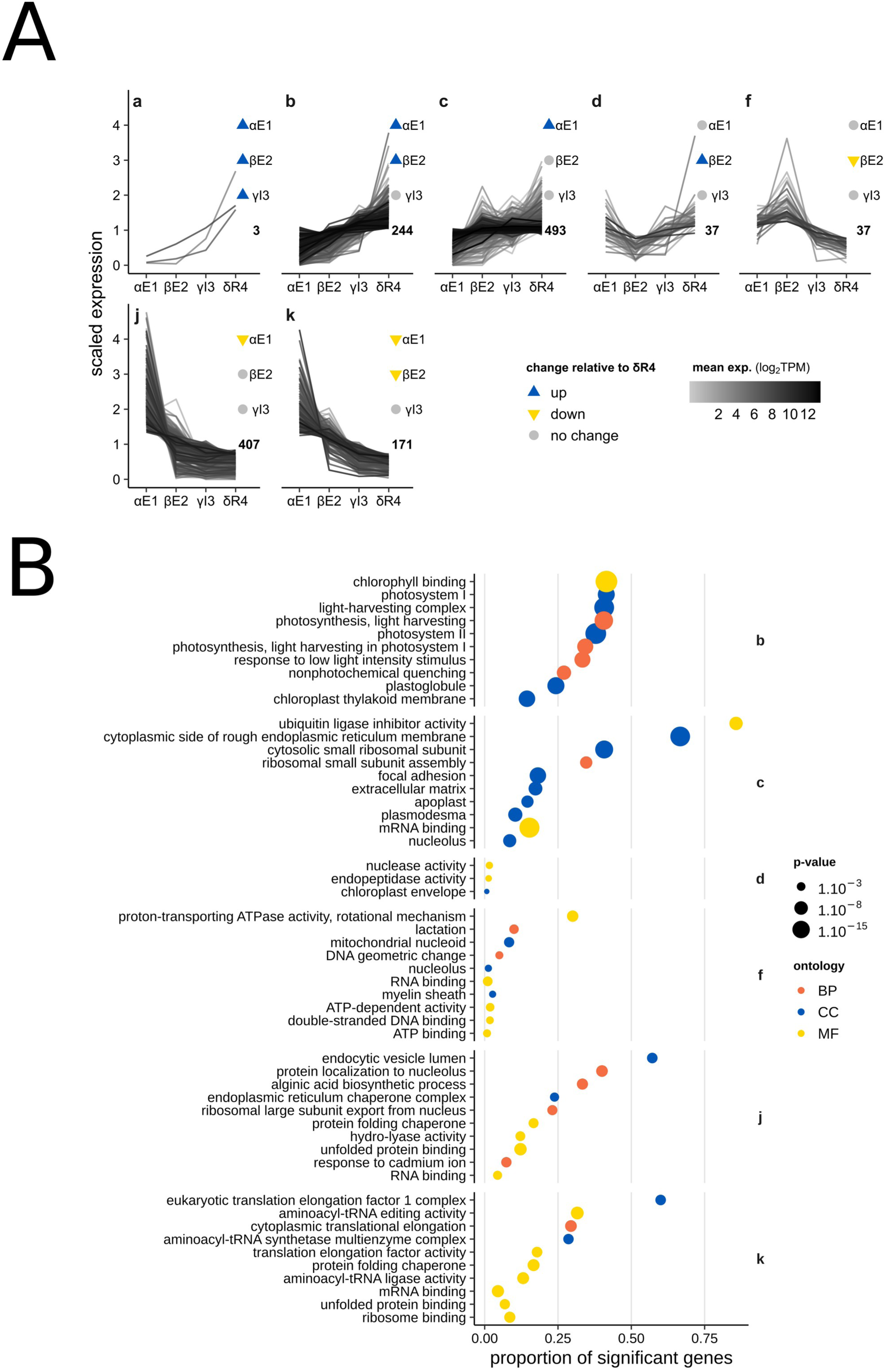
Clustering of genes significantly differentially expressed during αE1 to δR4 cell-type transition. **(A)** Gene expression in each cell type is represented as a scaled value of the gene expression (Transcripts Per Million; TPM) divided by the mean expression of the gene across all cell types. The mean expression in TPM (on log2 scale) of each gene is rendered on a black and white scale. Differently expressed genes were clustered based on the direction of change between either αE1, βE2 or γI3 in this order, and δR4. This algorithm was devised in order to capture genes that changed expression during the global transition toward the δR4 cell type, yet with too subtle changes between two adjacent cell types. Direction of change is indicated by blue (up) / yellow (down) triangles, or a grey dot (no significant change). The number of genes in each cluster is indicated in the lower right part of the graphs. Only clusters containing at least 3 genes are displayed. **(B)** Gene Ontology (GO) enrichment analysis of gene clusters during αE1 to δR4 cell-type transition. GO enrichment analysis was computed for clusters of differentially expressed genes composed of at least five genes using Fisher test and the elim algorithm for each ontology (BP: biological process; MF: molecular function; CC: cellular compartment). Terms represented by less than three significant genes were discarded. The proportion of significant genes (number of significantly differentially expressed genes divided by the number of annotated genes to a given GO term) was plotted only for the top 10 terms across the three ontologies. As a consequence, some clusters of differentially expressed genes are not represented here. Dot size is inversely proportional to the Fisher test p-value and coloured dots represent the term ontology.

We then derived gene expression profiles for each of these 6 clusters and performed GO term enrichment analysis (Fig. 2B;Table S3). The distinct identity of the apical cell was evident from the GO enrichment analysis of clusters ‘c’, ‘j’ and ‘k’. Despite having opposing trends in respect to the variation of transcript levels along the filament, all three displayed GO term signatures related to translation. This suggests that protein synthesis is paramount to the production of new cellular materials during growth of the apical cell and that the transition from αE1 to βE2 may involve a crucial change in the regulation of this process. Production of new cellular material is supported by the presence of the term ‘alginic acid biosynthetic process’ (Fig. 2B) and Golgi-related terms (Table S3) in cluster ‘j’. The former term is enriched because of higher expression in the apical cell of three different mannuronan C5-epimerase genes, two GDP-mannose dehydrogenase genes and one gene encoding a phosphomannomutase (Table S2). This is consistent with the combined biosynthesis of alginates, sulfated fucans and cellulose in the dome of the apical cell which is necessary for growth (Rabillé et al., 2019a; Rabillé et al., 2019b; Siméon et al., 2020).

Other clusters reflect additional functional features. Cluster ‘b’ comprises genes related to photosynthesis during differentiation from αE1 to δR4. This is consistent with the increase in photosynthetic pigments from the tip to the centre of the filament (Fig. S3), suggesting that the central cells supply apical cells with the carbohydrates necessary for tip growth. The enrichment of terms related to cell adhesion and communication in cluster ‘c’ may indicate an increased need for communication between filament cells, as well as for adherence to the substrate, during differentiation from βE2 to δR4.

We also compared the gene expression of branching cells (δB4) with that of primary filament cells. We hypothesised that these cells would display a transcriptomic profile similar to that of αE1 cells because branches grow by tip growth. The expression levels of 26 genes increased steadily from the αE1 cell to the δR4 cell before decreasing slightly in the δB4 cell (cluster ‘m’, Fig. S4A). GO enrichment analysis revealed that these genes were mostly related to photosynthesis (Fig. S4B). Conversely, the expression level of 31 genes decreased from αE1 to δR4 (cluster ‘n’) before increasing slightly in δB4. This result supports our hypothesis that δB4 cells resume the gene expression of αE1 cells. However, unlike in αE1, we did not identify any GO terms related to the biosynthesis of new material. Overall, our transcriptomics analyses indicate that the apical cell differs from the other cell types in the branched filament.

### Transcriptome signatures of cell types are lost in *etl* and *kna* mutants

To determine the extent to which cell shape, age and position influence cell-type specific transcriptome signatures, we examined the transcriptome profiles of cells in the *etl* and *kna* mutants and compared them with their equivalents in the WT. We selected cells based on their position in the filament: apical (cells αI2 and αE2 for *etl* and *kna,* respectively), sub-apical (cells βI3 and βE3 for *etl* and *kna,* respectively) and central (γδRB4,5 for *etl*, and γδE4,5 for *kna*; Fig. 1E).

First, we compared the overall transcriptome signatures of the mutant and wild-type (WT) in all cell types. A large proportion of genes of the WT transcriptome was expressed in the mutants: of the 12,999 gene transcripts expressed in the WT, 9,660 (74.3%) were also expressed in *etl,* 7,789 (59.9%) in *kna*, and 7,203 (53.8 %) were expressed in all three genotypes (Fig. 3). However, 2753 transcripts (20.6%) were expressed only in the WT, and 217 (1.6 %) and 94 (0.7 %) of genes were expressed specifically in the *etl* and *kna* mutants, respectively. This indicates a moderate but significant impact of the mutations on overall transcription.

**Figure 3:**
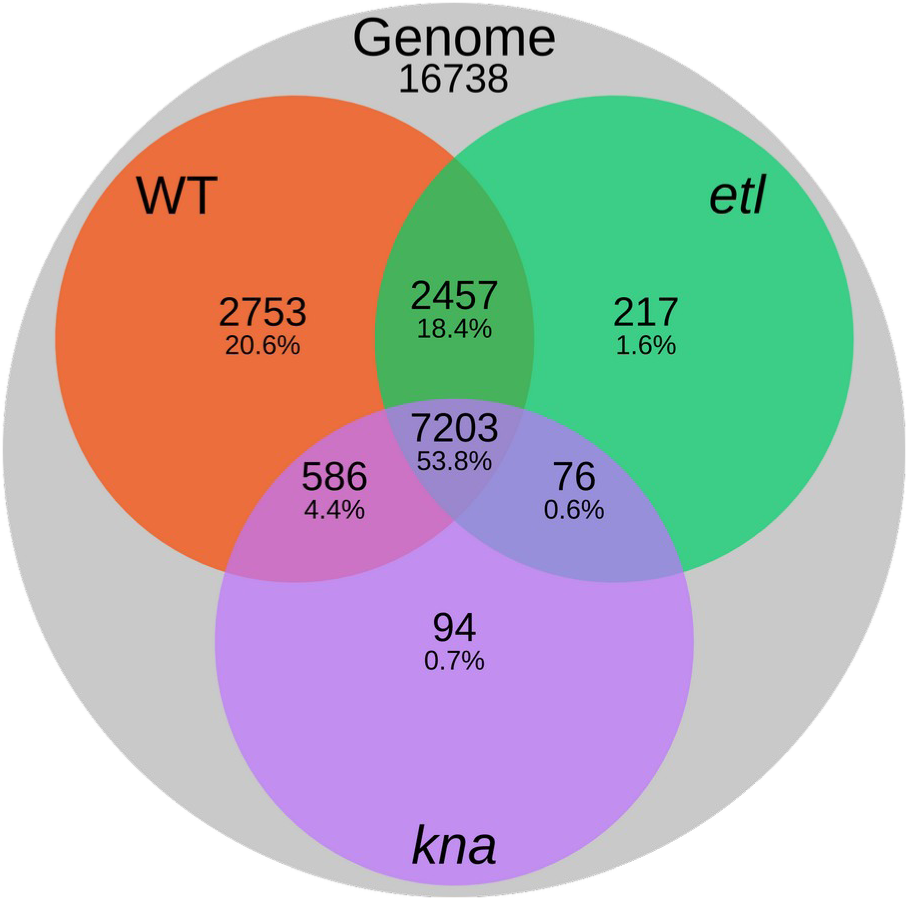
Venn diagram of the transcripts expressed in the WT, *etl* and *kna* genotypes. Genes with an average TPM value superior or equal to 1 in at least one of the five cell types were considered expressed in the corresponding genotype.

To understand better how altered cell age, shape and position affect gene expression in the mutants, we used principal component analysis (PCA) to analyse the transcriptomics dataset. To capture transcriptomic signals that might reflect differences between cell types and genotypes, we computed the ratio between the variance across conditions and the variance between replicates and performed the PCA with the 2500 genes with the highest variance ratio. When we analysed WT cells only, the samples clustered essentially by cell type, as expected (Fig. 4A). PC1 contained almost half of the variance in the dataset and mostly reflected cell position and shape along the filament, with αE1 on one side of the axis and δR4 on the other; βE2 and γI3 were located in between. The variance in PC2, with βE2 cells segregated from the other cell types, might be explained by a few genes with specific expression in βE2 cells, consistent with the clusters ‘d’ and ‘f’ shown in Fig. 2A. Interestingly, δB4 cells were located at an intermediary position along the PC1 axis, toward the α cell type, as expected since a new apex (α) emerges from these cells. This is consistent with clusters ‘m’ and ‘n’, as shown in Fig. S4, and is further supported by the PC3 axis, along which the δB4 cells are located away from the others (Fig. 4B). We assigned putative biological functions to the positive and negative sides of the PC3 axis by computing GO term enrichment based on the genes that correlated with them. We observed that the negative part of PC3 contained a high proportion of genes associated with intracellular signalling (e.g. response to virus, kinase activity) and cell division (Fig. 4C). This finding is consistent to some extent with the emergence of a new apex from the δB4 cell type and confirms that the branching cells are transitioning back to the α cell type through the activation of a few specific genes.

**Figure 4.**
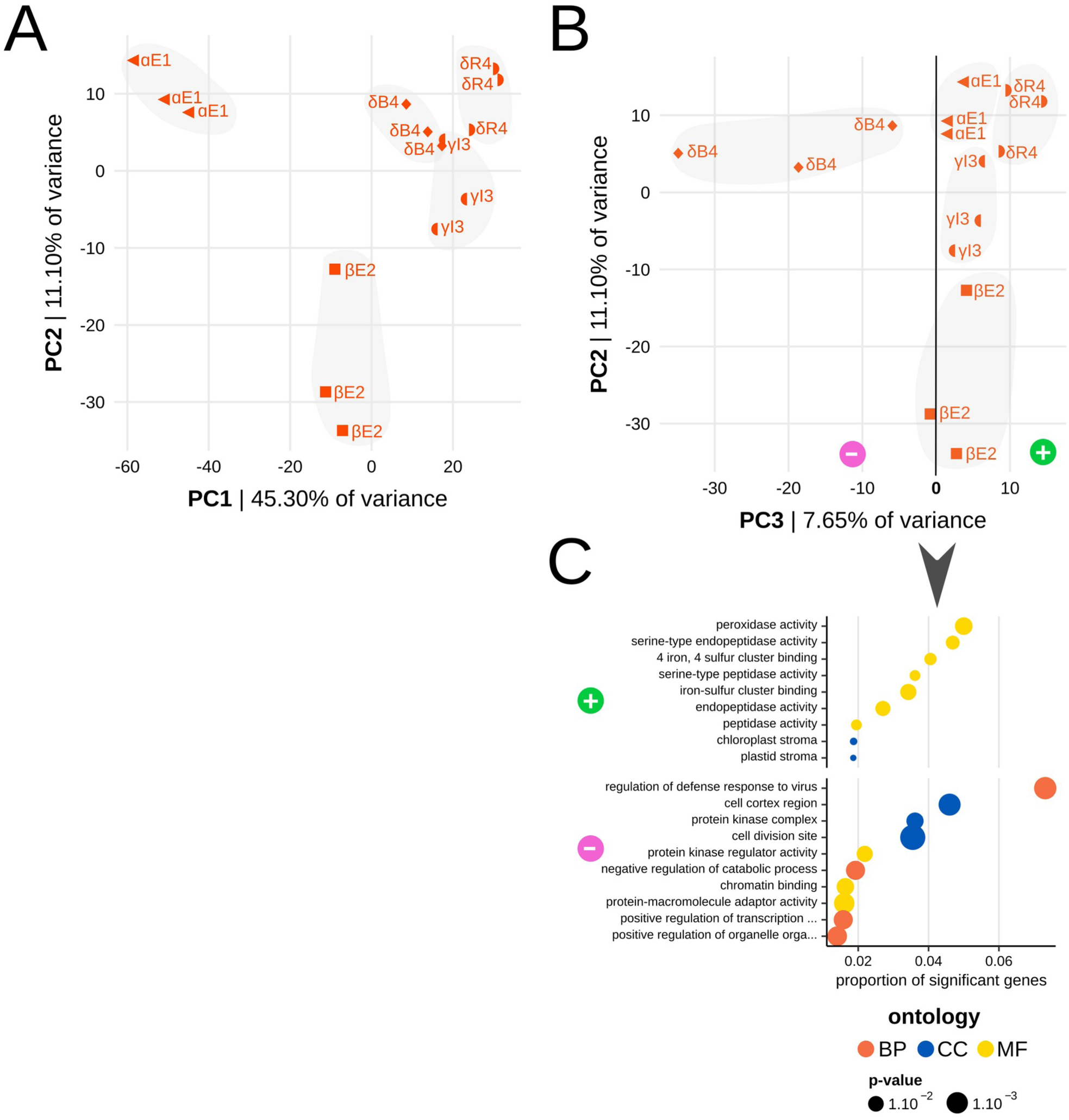
Principal Component Analysis of gene expression and GO term analysis in the WT. **(A, B)** PCA analysis with (A) PC1 and PC2 axes and (B) PC2 and PC3 axes. The first 2500 genes with highest VR were selected for PCA (see Material and Methods for explanation). Hand-drawn grey areas show clusters of samples of interest. **(C)** For the PC3 axis, genes that were significantly correlated (p ≤ 0.05) were obtained, and a GO term analysis was performed as shown in Fig. 2B, on the positive and negative correlations separately.

When we analysed the transcriptomics data from both the wild type (WT) and the two mutants using PCA, the samples clustered according to their genotype (Fig. 5A). The WT was found on the positive side of the PC1 axis and the two mutants on the negative side, irrespective of cell shape, age, or position (PC1 contained over half of the variance). This clustering likely reflects strong differences in gene expression between the three genotypes, as seen above (Fig. 3), which might be explained in part by the slower growth of the mutants when compared with the WT as shown in Fig. S2C. The positive side of the PC1 axis, where the WT cells clustered, was enriched in photosynthesis and translation terms (Fig. 5B; Table S3). The negative side, where the mutant cells clustered, was enriched with terms related to reverse transcription (RNA-directed DNA polymerase activity), the cytoskeleton (actin patches and filaments) and cell morphogenesis. This result is consistent with the two mutants being selected primarily for their altered cell morphology.

**Figure 5.**
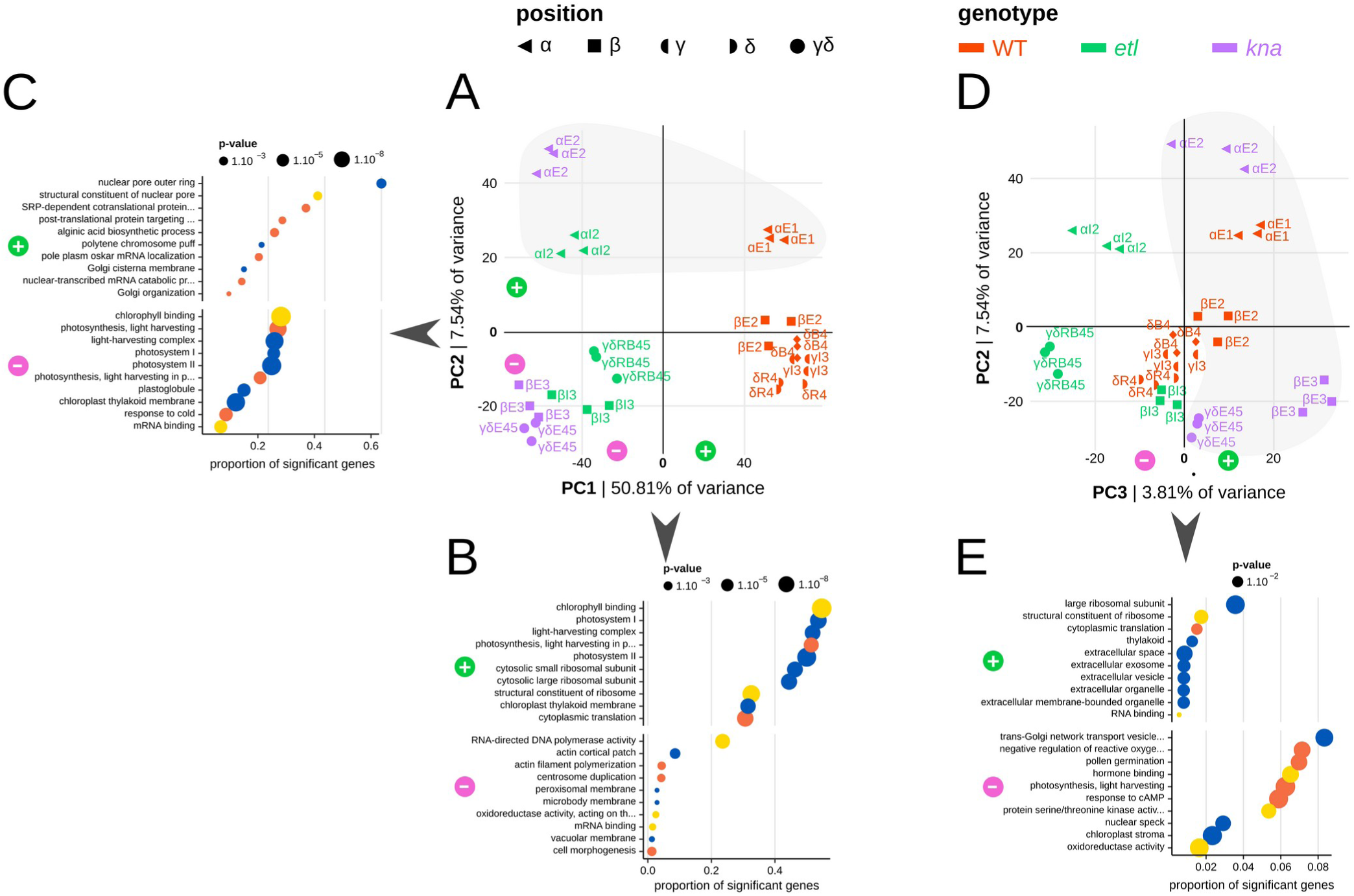
Principal Component Analysis of gene expression and GO term analysis in the WT and *Ectocarpus* mutants *etoile* and *knacki*. (A,D) Principal Component Analysis (PCA) of gene expression of the different cell types from the three genotypes WT, *etl* and *kna*. **(A)** PC1 and PC2. **(D)** PC2 and PC3. **(B,C,E)** GO term enrichment analysis of genes correlating to PCA axes. The positive and negative parts of each axis were analysed separately. **(B)** PC1 axis, **(C)** PC2 axis **(E)** and PC3 axis for all cell types.

On the PC2 axis (7.5 of the variance), cell types from the three genotypes clustered according to their relative position along the filament: α cells were all above the value 20 on the PC2 axis, whereas β, γ and δ position cells were below 0 and ordered from β to γδ coordinates (Fig. 5A). In *etl*, the γδRB4,5 cell type clustered between αI2 and βI3 cells, consistent with the star-like phenotype of this mutant, in which many α cells emerge from the center of the thallus, close to δ cells. In contrast to the correlation with cell position, neither cell shape nor cell age seemed to correlate with PC2. The elongated *kna* βE3 cells and the round WT δR4 cells, for example, had similar coordinates on the PC2 axis. A similar observation was made for young WT βE2 cells and old *etl* γδRB45 cells. While the α cells located in the positive part of PC2 display GO terms related to nuclear pores, RNA transcription and Golgi organisation, the cells located in the negative part of PC2 are involved in photosynthetic activities (Fig. 5C; Table S3). Based on the clustering along the PC2 axis, we conclude that cell position along the filament is the main factor determining cell identity.

Despite that PC3 represented only a small amount of the total gene expression variance (3.81%), it showed that all E cells (except one replicate sample of αE2) clustered on the positive side of the PC3 axis and displayed GO terms related to translation, exocytosis, and extracellular activities (Fig. 5D). By contrast, the other cell shapes clustered on the negative side of PC3 and displayed a variety of GO terms related to e.g. photosynthesis, the response to cAMP and hormone binding (Fig. 5E; Table S3). Based on this clustering along the PC3 axis, we conclude that the molecular identity of an *Ectocarpus* cell depends not only on its position along the filament, but also on its shape.

## Discussion

With its structure based on uniseriate filaments composed essentially of two morphologically distinguishable cell types and its relatively slow cell differentiation dynamics, *Ectocarpus* is an exceptional biological model for studying the relationship between cell shape, age and position within an organism. In the WT alga, apical growth results in the formation of elongated young cells at the apex, which transform into older, round cells in the centre of the filament. In the mutants *etl* and *kna* the relationship between cell position, shape and age is disrupted: in *etl*, apical growth rapidly arrests, resulting in round apical cells that cease to divide, whereas in *kna*, all cells, including those in the centre, are elongated. Both mutants grow more slowly than the WT. Using laser capture microdissection of the filaments of the WT and these two mutants, we find that cell identity, as defined by transcriptomics, depends primarily on the cell’s position along the filament, with cell shape contributing secondarily.

Nearly 80% of all genes in the genome are expressed in the early sporophyte phase of the life cycle. A significant proportion of these expressed genes were previously identified as being specific to the gametophyte. Such widespread genome expression in the sporophyte and gametophyte phases of the life cycle is typical of land plants and algae (Ratchinski et al., 2025; Somers and Nelms, 2023). Although this might be expected to hinder the identification of genes specific to a given cell type, we identified several hundred genes that are differentially expressed in adjacent cells within the *Ectocarpus* filament. Interestingly, variation in gene expression is not linear along the filament: there is a clear difference (472 differentially expressed genes) between the apical α cell and the subapical cells, suggesting that the apical position corresponds to a specific cell state. At those developmental stages, the apical cell is the only growing cell in the filament. This is consistent with its expression of genes involved in cell wall synthesis, the organisation of the Golgi apparatus, and the control of nuclear entry and exit — most likely of mRNA. However, we observed no significant enrichment of terms related to DNA replication itself.

How the apical cell might sense its unique position is a matter of conjecture. In fungi with hyphae, the elongated apical cells form an acid gradient from the apex to the centre of the filament (Limozin and Denet, 2000). In this case, the apical cell tends to accumulate protons because of active uptake and low ATP availability impairing proton outflow by proton pumps. In *Ectocarpus*, the apical cell has an extremely thin cell wall (Rabillé et al., 2019a), and exhibits low photosynthetic activity. It uses its energy to grow, making it poorly suited to expelling protons that enter through the thin tip. Thus, a proton gradient might be established along the apical cell of *Ectocarpus*, like that at the apex of fungal hyphae. This might result in the formation of an electrical field that serves as a spatial reference, as observed in the polarised zygote of the brown alga *Fucus* and also suggested during the embryogenesis and morphogenesis of other organisms (Nunes and Barriga, 2025).

Once the daughter cell of the apical α cell has formed, it loses its distal position and its expression profile changes radically from that of the apical cell. Then, it differentiates gradually as it reaches the centre of the filament by relative displacement. Only 218 genes are differentially expressed between subapical and median cells, and only 7 between median and central cells. Therefore, following apical cell division, the slope of cell differentiation is weak because all cell types display a highly similar molecular cell identity, as reflected by their expression profile. How cells know that they are no longer located at the apex of the filament, regardless of their shape and age, is as intriguing as the question of how they know that they are apical. Brown algae excrete organic matter, especially when they reach maturity (Halat et al., 2020; Powers et al., 2019). The sporophyte of *Ectocarpus* secretes diffusible signalling molecules (Arun et al., 2013) and inhibiting the vesicular trafficking using drugs blocks the development of the thallus (Green et al., 2013). Moreover, brown algae synthesise antimicrobial compounds (Lemesheva et al., 2023; Tajbakhsh et al., 2011) and the early sporophytes of *Ectocarpus* also seem to secrete components that inhibit the growth of bacteria (preliminary observations by B. Charrier and N. Serre). Hence, if all the cells of a developing filament (including the apical one) produce at the same rate a compound that does not diffuse rapidly, all the cells except the apical cell will be exposed to the same concentration of this compound. This self-organised spatial information may account for the cells’ perception that they are no longer apical, and result in major changes in their expression profile after the first, apical cell division, while normalising the expression profile of all the other, more central cells.

From an evolutionary perspective, it would be interesting to know whether filamentous organisms that are phylogenetically distant from brown algae have similar characteristics. Like the sporophyte of *Ectocarpus*, the gametophyte of the moss *Physcomitrium* develops a filamentous architecture through apical growth, composed of two main types of cells, chloronema and caulonema. These two cell types are distinguished by their growth rate, chloroplast abundance and cell division orientation (Coudert et al., 2019; Cove, 2005). However, the lack of distinct cell shapes precludes exploration of their shape–position relationship. Nevertheless, growth mutants of *Physcomitrium* (Peramuna et al., 2023) can be used to study the age-position relationship. (Xiao et al., 2011) identified approximately 400 differentially expressed genes in chloronema and caulonema, a similar number to those observed when we analysed the αE1 and δR4 cells of WT *Ectocarpus*. Unlike *Physcomitrium*, however, we found no GO terms associated with transcription factors, despite the expression of over 100 transcription factor genes in our dataset. Recently, (Chen et al., 2024) used enzyme-mediated cell dissociation and single cell RNA-Seq to characterise gene expression in the gametophyte of *Physcomitrium* during the transition from two-dimensional to three-dimensional growth. They found that the apical cells of these filaments preferentially expressed genes involved in photosynthesis and protein translation, while non-apical caulonemal and chloronemal cells were more involved in secondary metabolism, cell wall and membrane biogenesis. This contrasts with the situation in *Ectocarpus*, where photosynthetic functions are concentrated in non-apical cells, illustrating the diversity of cell differentiation strategies in filamentous organisms across the tree of life.

## Material and methods

### Wild type strains and cultivation

Two WT strains, male *Ectocarpus* species 7 (also named Ec32; CCAP 1310/4; origin San Juan de Marcona, Peru) and female *Ectocarpus* Ec568 strain (accession CCAP 1310/334; origin Arica, Chile) were used to perform crosses with the mutants. Thalli were grown in half-strength Provasoli-enriched (Starr and Zeikus, 1993), autoclaved natural seawater (NSWp; pH 7.8) in Petri dishes located in a controlled environment cabinet at 13°C with a 14:10 light: dark cycle (light intensity 29 mmol photon·m^−2^·s^−1^) as described in (Le Bail and Charrier, 2013). Light intensity and duration, temperature, external mechanical forces and gravity were maintained constant during the experiments.

### Mutants

The mutant *etoile* (*etl*; CCAP 1310/337) was generated from *Ectocarpus* species 7 as indicated in (Le Bail et al., 2011). It is a single locus mutant. In order to clear the genome from non causal mutations, *etl* was crossed with the WT female strain (Ec568) from which a female descendant was selected and back-crossed with the WT male *Ectocarpus* species 7 (Ec32). A female [*etl*] descendant of this second cross was used in this study. The mutant *knacki* (*kna*) was produced by UV-B irradiation of WT parthenogenetic gametes as reported in (Billoud et al., 2015; Le Bail and Charrier, 2013). It is also a single locus mutant. It was crossed with the female WT Ec568 and then back-crossed with the original WT male Ec32. A [*kna*] male descendant was selected for this transcriptomic study.

### Growth rate of Ectocarpus species 7 (Ec32), kna and etl

Single 2-week old *Ectocarpus* parthenosporophytes were first grown in 24-well plates in a drop of NSWp for 2 days to let them settle down. The wells were then filled with NSWp and pictures were taken every day for four days under the microscope Axio Observer.Z1 7 (Zeiss) with the objective N-Achroplan 5x/0.15 Ph1 M27. Growth rate of filament apices were manually measured using ImageJ (Schindelin et al., 2012) by taking branches as landmarks between each time point. N, number of measured filaments = 47 for WT, 39 for *kna* and 33 for *etl* genotypes.

### Age relative measurements

The age of cells can be defined based on two traits that occur simultaneously: First, the rate of cell division: the more a cell divides, the older its genomic DNA is considered to be, because of thereduction in the length of the telomeres. However, no study has measured the impact of the rate of cell division on the length of telomeres in brown algae. Therefore, we considered the second trait to be more relevant: the rate of biosynthesis of new material. The faster a cell grows, the more recently its material was formed. In the WT, the apical cell grows rapidly, resulting in the cell being composed of new material. We assigned it the arbitrary number 1. The mutants grow more slowly than the WT, so the material composing their apical cell is older. We assigned the apical cells of these mutants the value 2. In both the WT and the mutants, all the other cells are produced successively by the apical cell. As a result, their ages are identified by the following numbers: 2–4 for the WT and 3–5 for the mutants. See Fig. S2D for a detailed explanation.

### Laser capture microdissection

Germination of *Ectocarpus* species 7 WT and mutants took place directly on PEN slides (ThermoFisher Scientific LCM0522), as described in (Saint-Marcoux et al., 2015). Three slides were simultaneously cultivated in separate Petri dishes for each of the WT and the two mutant strains, and were considered as independent replicates. Fixation was performed 10 days after germination. Cells of filaments of 15-20 cells were preferentially captured. WT series #2 was fixed 5 days after WT series #1 and #3. The mutants and the wild type (WT) were captured in separate experiments.

### RNA extraction

PEN slides with ∼ 30-cell filaments were fixed and tissue extracted by laser capture microdissection as described in (Saint-Marcoux et al., 2015). RNAs were extracted and amplified as described in (Saint-Marcoux et al., 2015). Filaments were cut in four regions labelled α, β, γ and δ from the apex (position α, representing one single cell) to the center of the filament (position δ). Apart from the α region, all the other regions of a single filament contains 2-4 adjacent cells. Mutant cell types followed the same experimental procedures as the WT. However, because of their morphology, — bushy for *etl*, and hypo-branched for *kna —* or for the lack of clear cell differentiation based on cell morphology, only three regions were dissected, named α, β, and γδ (Fig. 1E). Branching cells were also captured separately from the WT organims. In *etl*, branching cells were mixed with the central domain γδ and in *kna*, they were absent.

### RNA-seq analysis

For each sample, 10 Millions of Paired-End reads were produced with the HiSeq NGS technology (BGI, Table 1 in Supplementary information). The RNA-seq reads were filtered using Trimmomatic (Bolger et al., 2014) and rRNAs were removed by SortMeRNA (Kopylova et al., 2012). The remaining reads were mapped onto the *Ectocarpus* genome and transcriptome V2 representing 18271 genes (Cormier et al., 2017) available in Orcae (Sterck et al., 2012) using bowtie (Langmead et al., 2009) and RSEM (Li and Dewey, 2011). Further analyses were performed only on genes not involved in sexual differentiation (Lipinska et al., 2015) since the WT and *kna* were male and *etl* female, thus leaving 16738 nuclear genes. Raw gene expression was length corrected with tximport (Soneson et al., 2016) and DESeq2 (Love et al., 2014) was used to obtain differentially expressed genes in the WT (adjusted p-value for multitesting ≤ 0.05). The ashr algorithm was used for log fold change shrinkage. Variance stabilising transform expression matrix was obtained on the entire dataset containing the gene expression of the three genotypes. Gene clustering was conducted by defining patterns of expression after differentially expressed genes from different pairwise comparisons as described in the legend of Fig. 2A. GO enrichment analyses were performed using topGO package v2.58 after *Ectocarpus* genome annotation was augmented by scanning the predicted peptides against the Swiss-Prot database release 2022_01 using Blast, and the Hectar pipeline to predict transit peptides (Gschloessl et al., 2008). All statistical analyses and graphs were obtained using R v4.4.3 (R Core Team, 2025). The fluctuations in expression levels observed by NGS were confirmed by Q-PCR on a partial sample (Supplementary Information).

### Selection of genes for the Principal Component Analysis

The variance ratio of each gene was computed as follows: 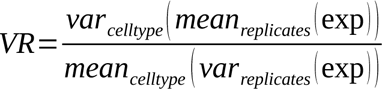 with exp being the expression level of the gene. The 2500 genes with the smallest variance between replicates and highest variance between cell types were selected for the PCA.

## Acknowledgement

Research was funded by PEPS “ECTODIFF” from C.N.R.S. (B.C.) and the European Research Council (project ERC-Adv EDIP, grant number 233185; J.A.L.). The authors are thankful to Elodie Rolland for the maintenance of the algal cultures.

## Authors contribution

D.S.M and B.B. performed the sequence and statistical analyses, C.D. and B.B. performed the time-lapse monitoring of cell rounding. S.C. performed the Q-PCR. B.C. and D.S.M. performed laser capture microdissection. D.S.M. performed the RNA extraction and amplification. B.C. designed the biological experiments and wrote the initial versions of the paper. D.S.M, B.B, J.A.L and B.C contributed to the final version of the manuscript.

For the purpose of Open Access, a CC-BY-NC public copyright license has been applied by the authors to the present document and will be applied to all subsequent versions up to the Author Accepted Manuscript arising from this submission.

---

**Movie 1: Time-lapse of *Ectocarpus* growing sporophyte.**

EctocarpusDevelopment_01.webm shows a time-lapse movie of a young WT *Ectocarpus* sp. sporophyte. αE1 (apical, elongated and young) cells grow and periodically divide to give βE2 (new sub-apical elongated) cells. These βE2 cells then get rounder, passing through the γI3 (proximal, oval old cell) stage before they eventually become δR4 (centered, round very oldest cells). Occasionally, cells develop lateral branches: they are called Branched (δB4) and give rise to a novel filament led by an αE1 cell. Time is given in hours.

## Supplementary figures

**Figure S1:**
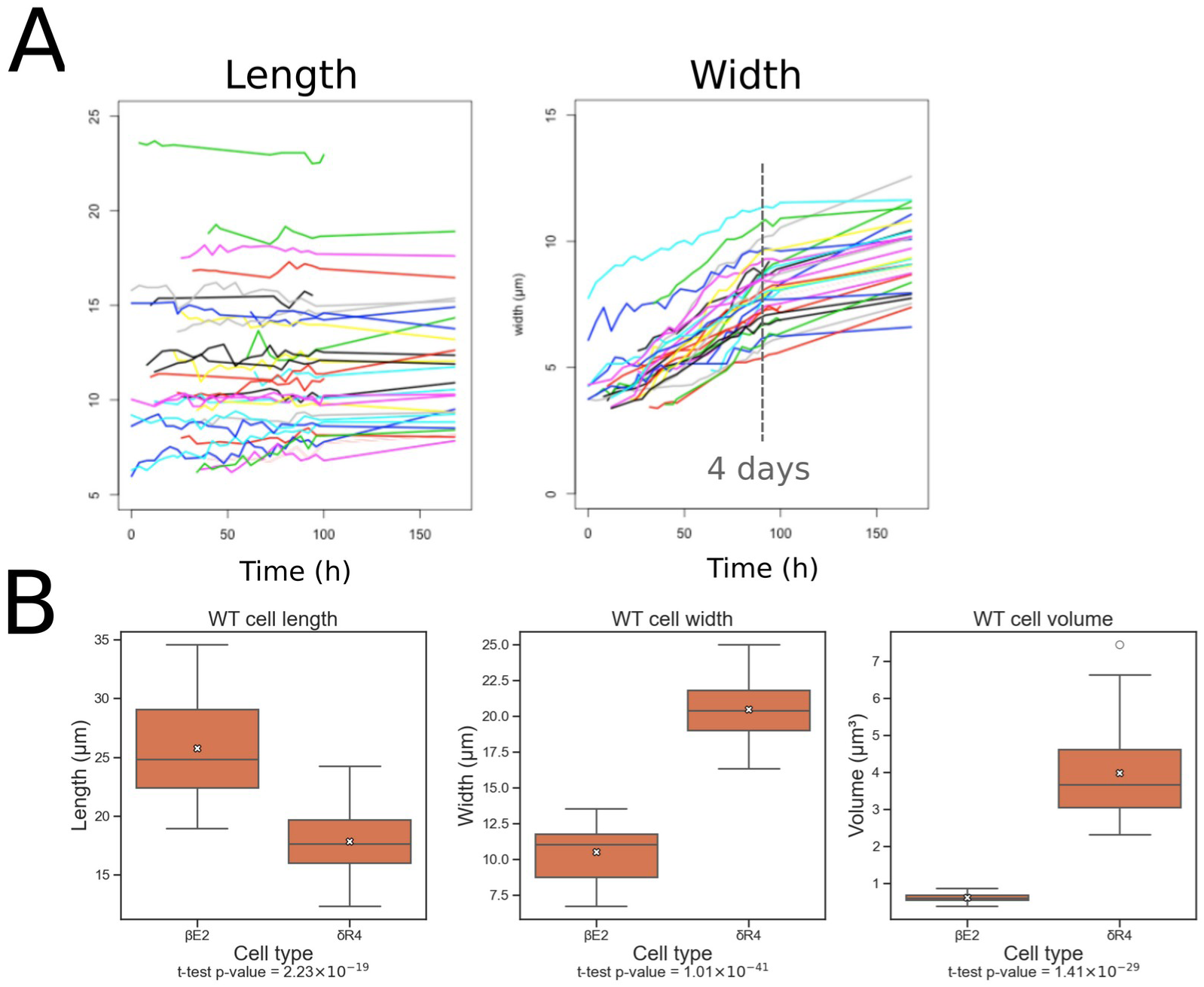
Cell morphometrics and differentiation rate in the sporophytic filament of WT *Ectocarpus sp*. **(A)** Rate of cell rounding. Cell length (i.e. along the filament axis; left) and width (i.e. perpendicular to the main filament axis; right) of *Ectocarpus* WT βE2 were measured every 2 hours over 7 days. Each colour represents a βE2 cell from a different individual. **(B)** Quantification of the length, width and volume of *Ectocarpus* WT βE2 and δR4 cells. The length (µm; left), width (µm; middle) and volume (µm^3^; right) of βE2 (blue) and δR4 (pink) cells were measured. Width was measured at the centre (middle right) of the cell considered along its longest axis. Cell volume was calculated from the cell length and width considering the cell as a regular ellipsoid for δR4 cells, as a truncated ellipsoid (barrel) for βE2 cells. Median is shown with the horizontal bar, mean with the white cross, the box shows the 2^nd^ and 3^rd^ quartiles, the whiskers extend to the most extreme values within the box, which is ±1.5 times the interquartile range. Results of Student’s t-test are indicated below each plot.

**Figure S2:**
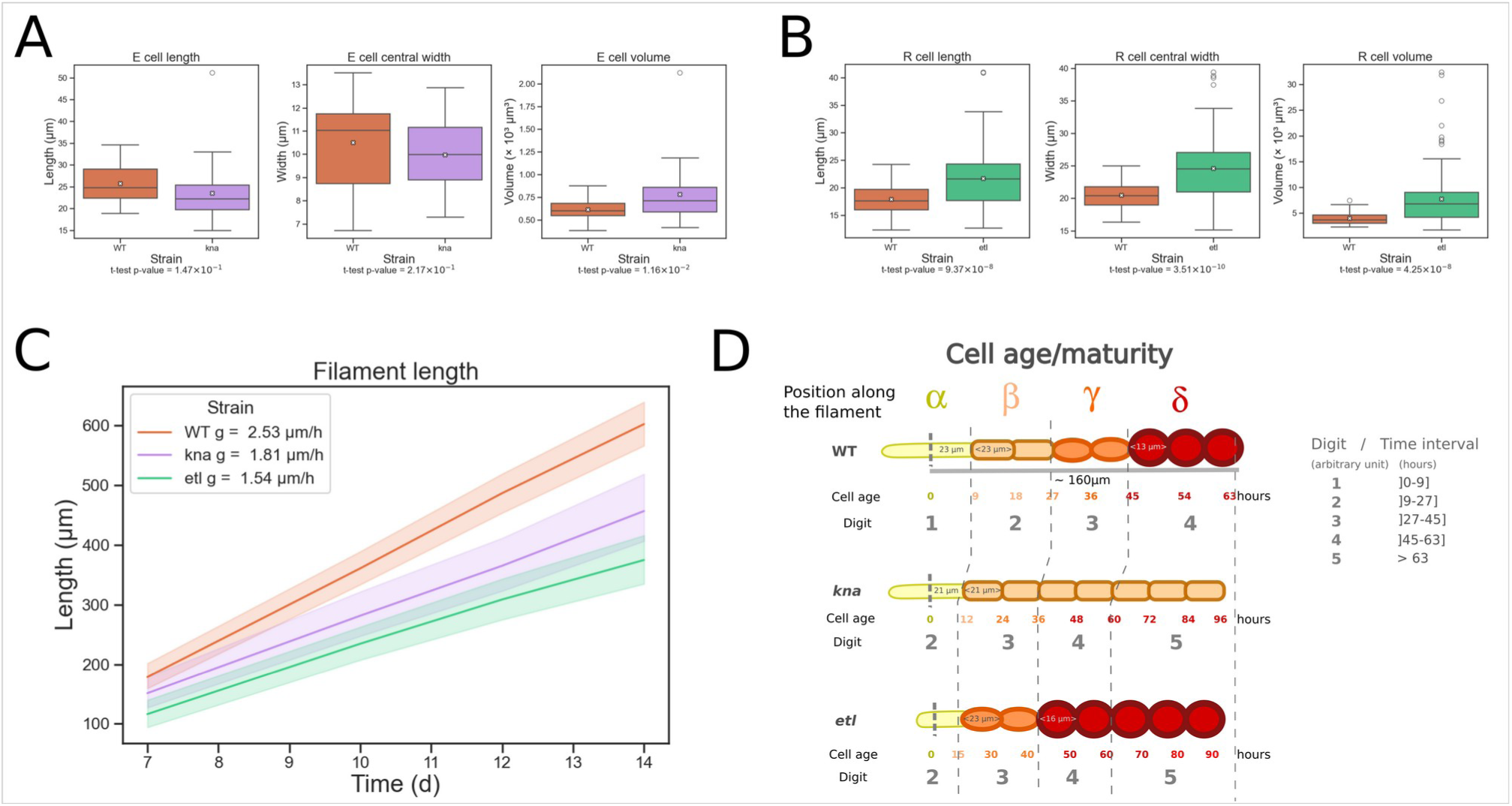
Cell morphometrics and growth dynamics for the mutants *kna* and *etl*. **(A,B)**: For each mutant (**A:** *kna;* **B**: *etl*), the length (left panel), width at the cell center (middle panel) and cell volume (right panel) were measured and compared to those of the WT (orange boxes). Cell volume was calculated as in Fig. S1B; box-plot was drawn as in Fig. S1B. **(C)** Growth rate (g) in the WT, *kna* and *etl* mutants. The increase in filament length (µm) was measured over the course of time (days ‘d’). n>50 replicates. Slopes of the growth rate curve is indicated for each genotype in the insert in the top left of the graphic. **(D)** Relative cell age. Relative cell age was calculated for each cell of a half filament of 8 cells (total filament size would be ∼ 16 cells), which measures about 160µm in the WT. This stage corresponds to the filaments that were microdissected in the WT and in the mutants. Cell shape and size (based on literature; Le Bail et al., 2008; Le Bail et al., 2011) are indicated on the schematics with a colour code (yellow: alpha; orange: beta; dark orange: gamma; dark red: delta). Cell age (expressed in hours, first row below the schematics) was calculated from cell position along the filament, taking into account the cell size and the measured growth rate shown in Fig. S2C. The digit given to each position (second row below the schematics) represents the approximate cell age, based on the dynamics of cell differentiation and growth in the WT. Digit 1 for cells comprised between [0;9] hours; digit 2 for [9;27]; digit 3 for [27; 45] and digit 4 for cells older than 45 hours. The reference time “0” corresponds to the moment when the apical cell of each genotype divides. This takes 9 hours in the WT, 12 hours in the *kna* mutant, and 15 hours in the *etl* mutant.

**Figure S3:**
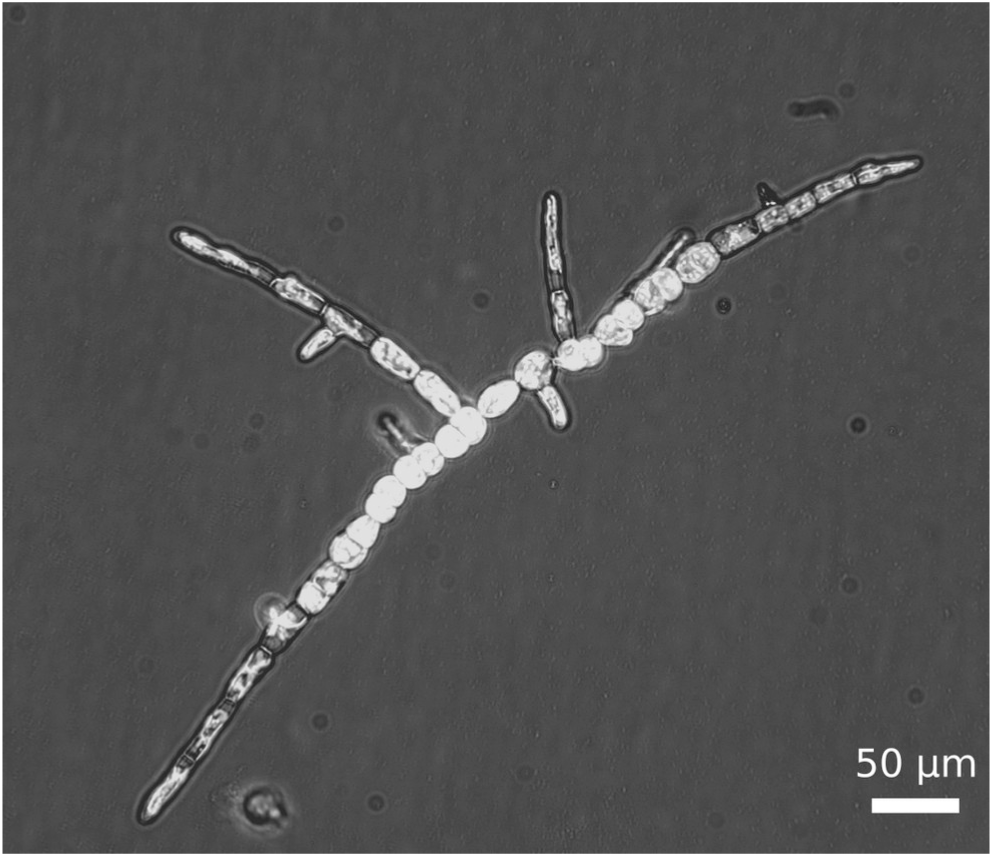
Autofluorescence of a branched filament of an *Ectocarpus* sp. sporophyte. The white areas correspond to the autofluorescence of the chloroplasts that were excited using a UV lamp on a DMI8 inverted microscope (Leica Microsystems).

**Figure S4:**
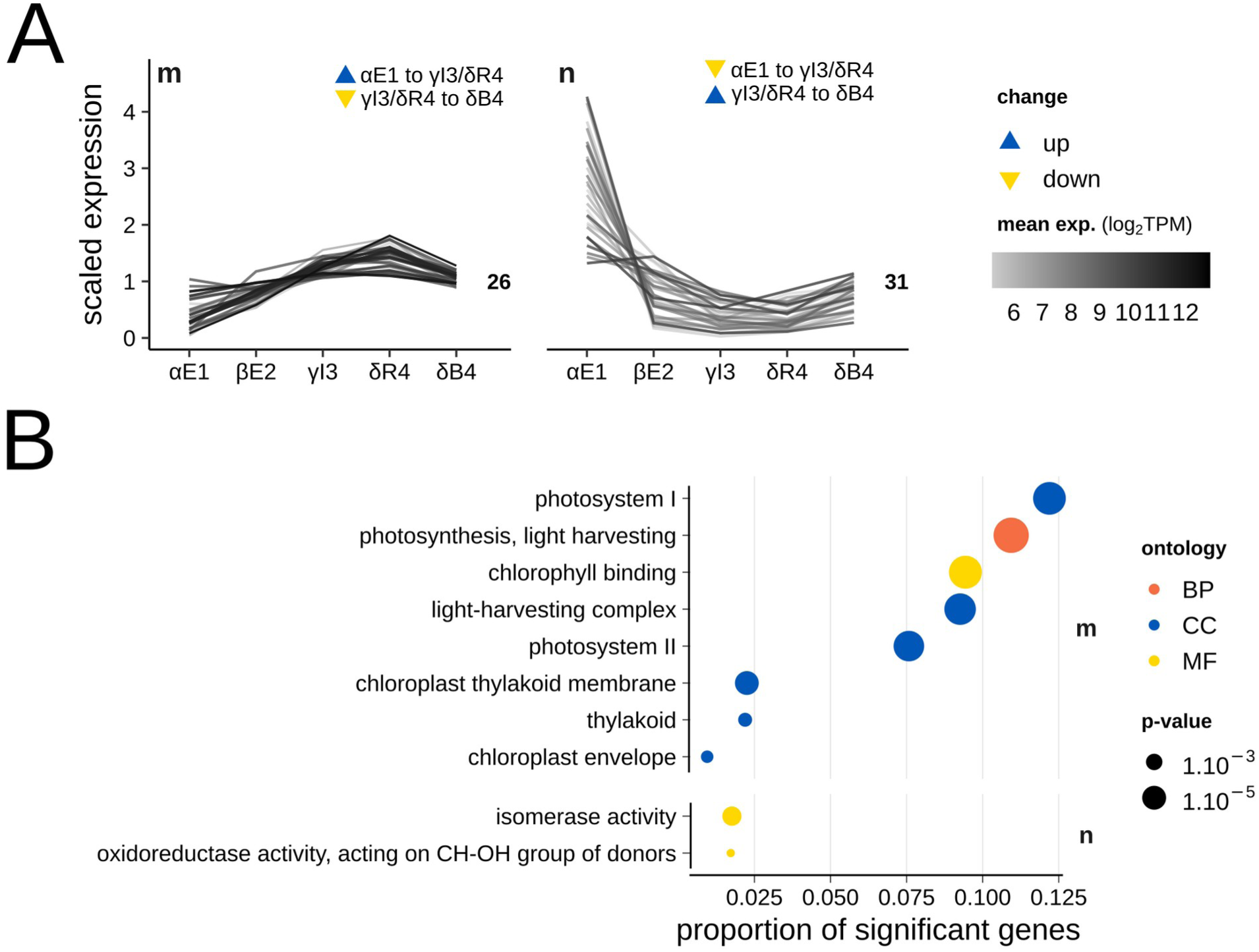
Expression profiles and GO enrichment analysis of genes potentially related to branching. **(A)** As in Fig. 2A, genes were clustered in the same group when the direction of change between αE1 and γI3 or δR4 was opposite to that between γI3 or δR4 and δB4. The reasoning was that genes defining αE1 cellular status would have a similar expression profile in the δB4 cell type, and an opposition profile in δR4 cells. **(B)** GO term analysis, as in Fig. 2B.

## Supplementary tables (attached zip.file “Table_Sn.zip”)

**Table S1.** Sporophytic- and gametophytic-biased and specific genes expressed in the early stage sporophyte.

**Table S2.** Differentially expressed genes between cell types in WT, and in each mutant *etl* and *kna*.

**Table S3.** GO-term enrichment in lists of differentially expressed genes between cell types in WT, and in each mutant *etl* and *kna*.

**Supplementary information** (attached zip file “Supplementary_Information.zip”)

