## Supplementary figures and images for "Cell position is the primary determinant of cell identity in the uniseriate filament of the brown alga *Ectocarpus*"

### Suppl_Fig_1_Supplementary_Information.png

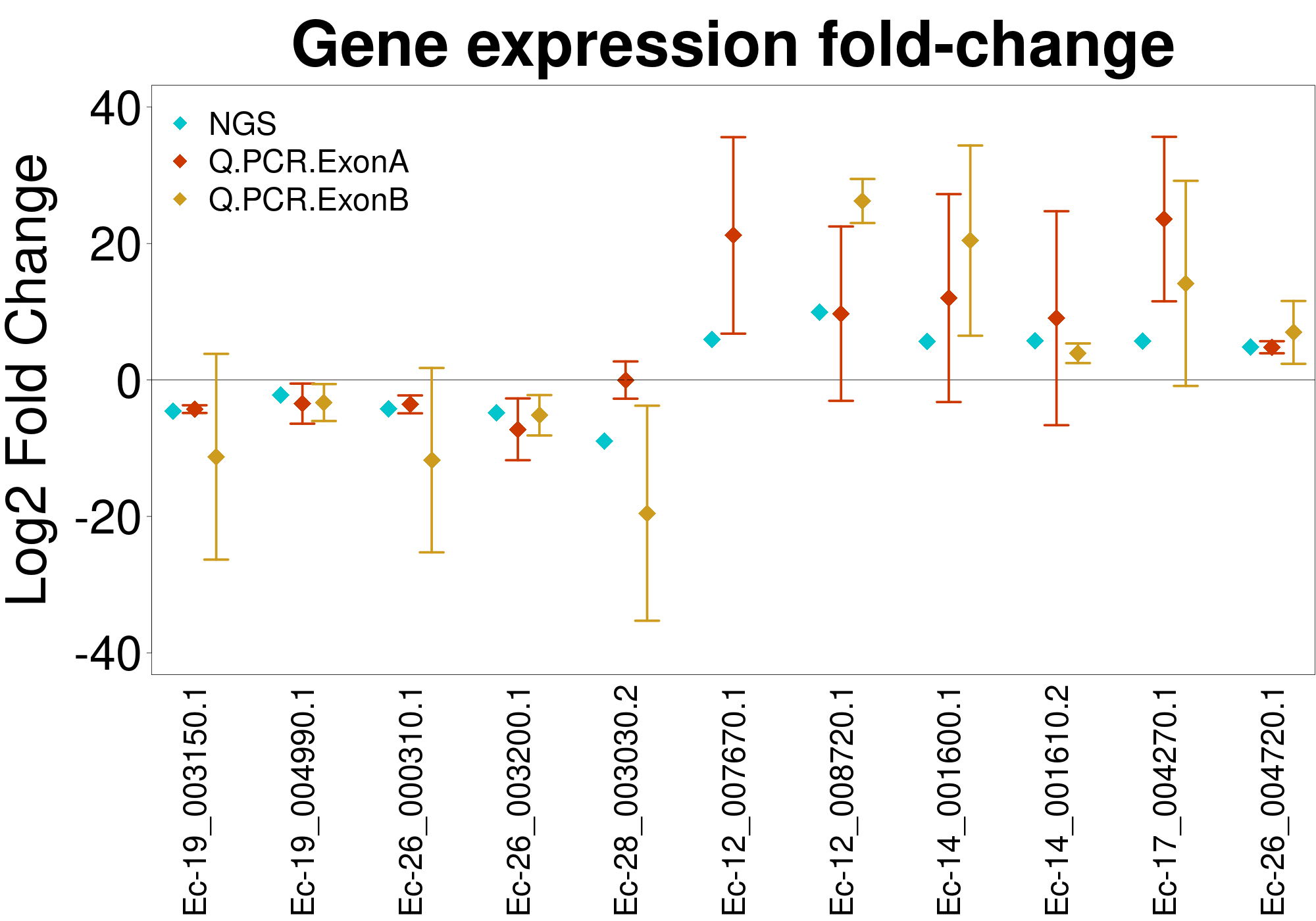
